# Low effective population size but high heterozygosity revealed by SNP analyses in adult and juvenile *Thuja plicata* in UK Woodlands

**DOI:** 10.1101/2024.03.18.585477

**Authors:** Laura Guillardín, Ella Glover, Gary Kerr, John J. MacKay

## Abstract

*Thuja plicata* is a conifer tree that is appreciated for its cultural, ecological and wood quality features in its natural range in western North America. It is also used in Europe and the UK for timber production. Some *T. Plicata* plantations in the UK are converted to Continuous Cover Forestry (CCF) management which uses natural regeneration and concerns have been raised about the genetic diversity in both the adult trees and the offspring. We studied species diversity, stand structure, and genetic diversity in four UK woodlands planted with *T. plicata* which contained both adults and naturally regenerated individuals. We discovered 61,104 Single Nucleotide Polymorphisms (SNPs) using Genotyping-by-Sequencing (GBS) and retained 504 SNPs for analysis. We selected 67 SNPs for PCR-based genotyping of 163 adults and 176 juveniles. We found a large number of monomorphic sites (40.3%) most of which were restricted to adults in a single woodland, indicative of a high genetic differentiation among woodlands. The effective population size (N_e_) was very low across all sites (N_e_ < 100), varied between adults and juveniles, and was lowest in the most diverse woodland in terms of species and structure. In contrast, heterozygosity was high overall (H_o_ = 0.38), except in the divergent woodland (H_o_ = 0.26), and did not vary between adults and juveniles. Our findings and genotyping methods provide insights into the ability of CCF-managed woodlands, including instances of low genetic diversity in *T. plicata*, to face environmental shifts and disease threats.

## Introduction

*Thuja plicata* Donn ex D.Don is a large evergreen coniferous tree native to western North America, which may be found in UK woodlands under Continuous Cover Forest (CCF) management. The CCF management approach uses a variety of practices adapted to local conditions including the use of natural regeneration to renew forest stands. It ultimately aims to increase and safeguard biodiversity while preserving the timber harvest. CCF is viewed as more effective at curbing the impact of climate change than forest management practices based on clear-felling (Helliwell and Wilson 2012; Mason et al. 1999) and as suitable for extending the range of ecosystem services that are provided to the community (Kirby and Watkins 2015). The stand structure is defined by the spatial distribution, species diversity and variation in tree dimensions (Pommerening 2002). However, the genetic makeup of the woodlands may be altered by CCF, with a potential decrease observed in the genetic diversity over generations (Macdonald et al. 2010; Finkeldey 2002). The use of plantations of exotic species such as *Thuja plicata* in the development of CCF raises further questions as to their diversity and ultimately their potential adaptability to changing environmental conditions.

*Thuja plicata* has been used and managed by First People Nations in Northern Western America for centuries for cultural and utility items (Hebda and Mathewes 1984). Due to its valuable wood characteristics such as durability and resistance to rot (Stirling et al. 2017), *T. plicata* is recognised as producing high quality timber for construction and is widely planted in Europe for this very reason. *T. plicata* gained importance in UK forestry in the late eighteenth century (Jarvis 1973) but information on the origin and genetic makeup of UK plantations is either lacking or scarce.

*T. plicata* is monoecious, i.e. individuals have both the male and female reproductive structures as separate organs, and anemochorous, i.e. their pollen and seed are wind-dispersed. Low genetic diversity levels have been found in its natural range based on heterozygosity (H_e_ and H_o_), nucleotide diversity (π) and effective population size (N_e_) (Shalev et al. 2022), which could potentially impact the species’ long-term adaptability. While *T. plicata* is principally outcrossing, there are reports of inbreeding by mating genetically related individuals or self-reproduction in some situations. Although *T. plicata* has relatively high selfing rates, it has low levels of inbreeding depression in growth and wood chemistry traits (Shalev et al. 2023; Wang and Russel 2006). Genetic diversity varies along a latitudinal gradient within its natural distribution, showing higher levels in the south coastal region (O’Connell et al. 2008). This southern region was a refugium for *T. plicata* during the last glacial period and it later spread to the north and east to its current natural distribution (O’Connell et al. 2008). Predictions suggest potential further spread from coastal areas towards the interior due to changing climatic conditions (Hamann and Wang 2006). However, climate change may surpass the potential for adaptation in several species including *T. plicata* (Alberto et al. 2013), which could affect populations in both the natural range and in areas of artificial introduction.

There are a few genomic resources to support investigations in *T. plicata* including a transcriptome (Shalev et al. 2018) and a first reference genome (Shalev et al. 2022). Restriction site-associated DNA sequencing (RADseq) and Genotyping-by-Sequencing (GBS) have emerged as powerful and cost-effective tools for genotyping in non-model organisms (Holliday et al. 2018). GBS was originally described as a method using DNA digestion with a single restriction enzyme (Elshire et al. 2011), while double digestion RADseq (ddRADseq) used both common and rare cut-site enzymes to optimise fragment sizes and achieve the desired genome coverage (Parchman et al. 2018). Nowadays, GBS is used to describe methods for sequencing and genotyping which use one or several restriction enzyme cutting sites (Andrews et al. 2016). GBS is useful allows the direct discovery of thousands of single nucleotide polymorphisms (SNP) without prior information; however, GBS requires deep data curation and complex bioinformatic pipelines (Vaux et al. 2023, Guillardín-Calvo et al. 2019). For example, estimating genotyping errors (Mastretta-Yanes et al. 2015) and missing data handling by imputation (Mora-Márquez et al. 2023) are key steps to process GBS data to obtain reliable results. Alternatively, when prior information is available on sequence variation in the populations of interest and a reduced number of markers is sufficient, other genotyping methods are available that may be simpler to analyse, have less missing data, increased reliability, and may reduce costs. High-throughput platforms including Illumina Infinium iSelect HD (Illumina, San Diego, CA), the Sequenom MassARRAY platform (Agena Bioscience, San Diego, CA), and Standard Biotools Dynamic Arrays (San Francisco, CA, USA) offer a range of genotyping approaches suitable for large numbers of samples that are cost and time-effective (Choi et al. 2020; Davey et al. 2011).

Our hypothesis is that materials used to plant *T. plicata* woodlands being converted to a CCF may have low standing genetic diversity. In addition, selective logging of individual trees may decrease the effective population size resulting in naturally-regenerated juveniles with a reduced genetic diversity relative to initial plantations. We tested this hypothesis by analysing planted *T. plicata* trees from four different woodland sites under CCF management in the UK. We first used GBS for SNP discovery with two enzymes to digest the *T. plicata* DNA and selected a subset of informative SNP markers for population-scale genotyping. Our objectives were: 1) To characterise the woodland level of transformation from plantations toward CCF; 2) To perform SNP discovery in *T. plicata*; 3) To design and validate a reduced SNP genotyping set; 4) To assess the genetic diversity differences between adults and juveniles across sites.

## Material and Methods

### 1. Study sites characterisation: structure and species diversity

We sampled four CCF sites located in England (UK) that included *T. plicata* in the canopy, identified as Bagley Wood, Longleat1, Stourhead1, and Stourhead2 (Table 1). Each site had at least 100 adult trees as recommended by Wojacki et al. (2019) to avoid biases from sampling small populations, as well as having enough viable natural regeneration. We quantified the total surface area of each site using QGIS software (QGIS 2023) and defined the stand distribution as patchy if the site includes separated areas, or as continuous (Table 1). We assessed the relative isolation sites by determining the presence of *T. plicata* woodlands within a 2 Km radius from their center, based on the National Forest Inventory Woodland public dataset (Forestry Commission 2019) and private land cover shapefiles (supplied by landowners).

**Table 1.**
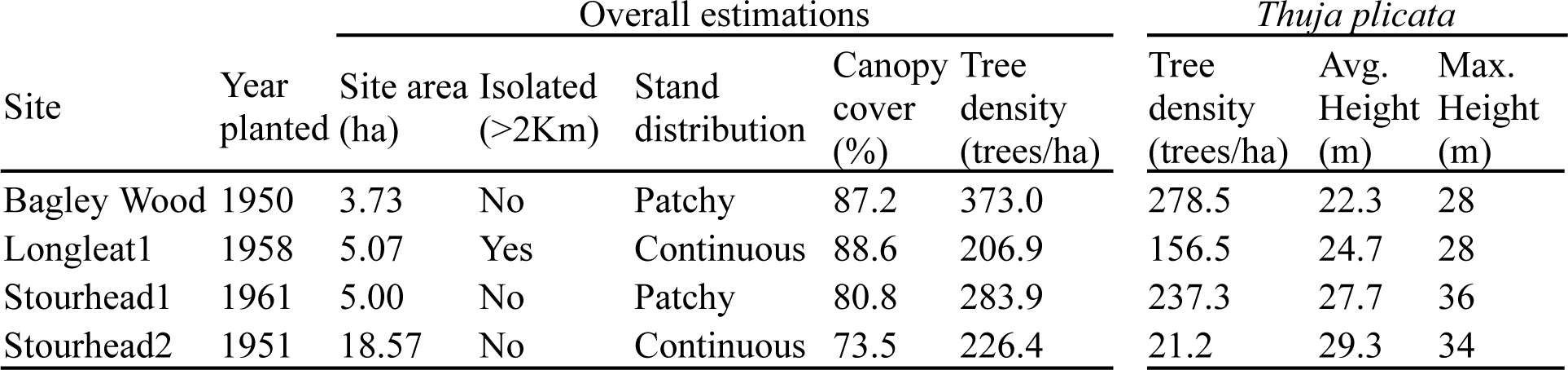
Site characterisation results, including overall site estimations and specific estimations for *Thuja plicata* trees.

| Site | Year planted | Overall estimations |  |  |  |  | *Thuja plicata* |  |  |
| --- | --- | --- | --- | --- | --- | --- | --- | --- | --- |
|  |  | Site area (ha) | Isolated (>2Km) | Stand distribution | Canopy cover (%) | Tree density (trees/ha) | Tree density (trees/ha) | Avg. Height (m) | Max. Height (m) |
| Bagley Wood | 1950 | 3.73 | No | Patchy | 87.2 | 373.0 | 278.5 | 22.3 | 28 |
| Longleat1 | 1958 | 5.07 | Yes | Continuous | 88.6 | 206.9 | 156.5 | 24.7 | 28 |
| Stourhead1 | 1961 | 5.00 | No | Patchy | 80.8 | 283.9 | 237.3 | 27.7 | 36 |
| Stourhead2 | 1951 | 18.57 | No | Continuous | 73.5 | 226.4 | 21.2 | 29.3 | 34 |

We conducted a forest survey to estimate the degree of transformation from a plantation towards a CCF woodland (Figure 1) following the methodology recommended in the Forest Inventory and monitoring guidelines (Northwest Natural Resource Group 2014). We used QGIS (QGIS 2023) to randomly select the centre of up to 25 plots on each site. The radius of plots ranged from 6m to 8m, depending on the site, and the natural regeneration plots were 3m across (Figure 1). We measured diameter at breast height (dbh) using a diameter tape, and height using the Bitterlich Relascope method (Bitterlich 1984) on every tree inside the plot. We measured the dimensions of regenerated seedlings and saplings within the regeneration plot using a digital calliper and a metre stick and took a census of each species. We estimated the canopy density of each plot by using a densiometer (Figure 1) and the *T. plicata* tree density of each site.

**Figure 1.**
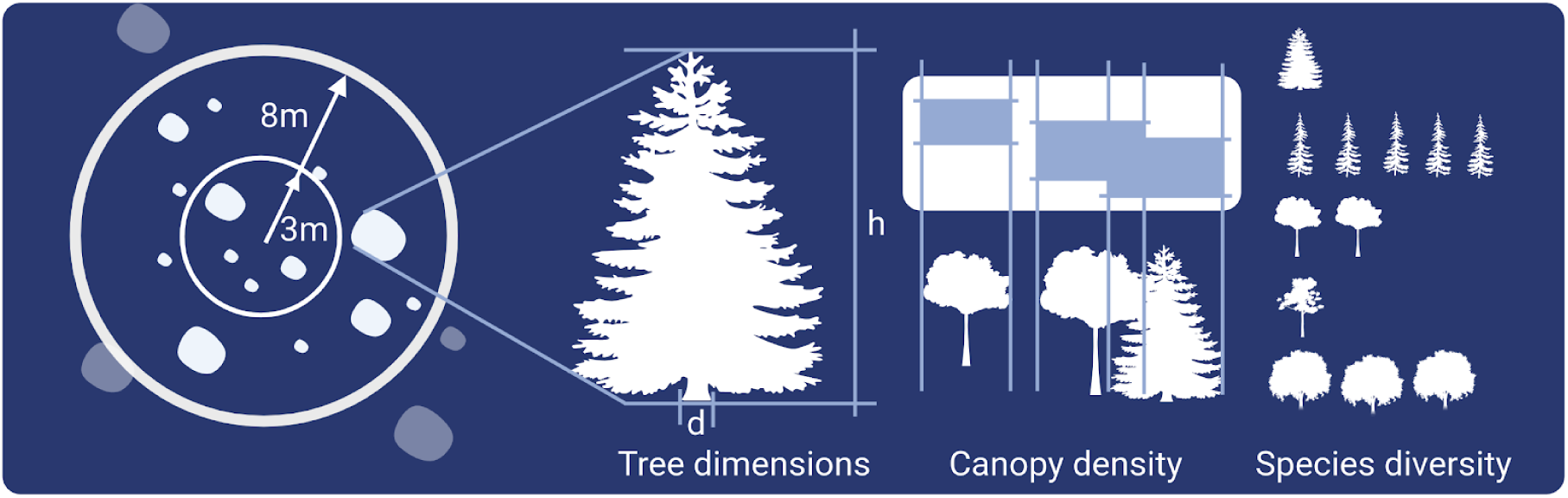
Forest inventory plots description and parameters measured. Tree dimensions: d = diameter at breast high, h= tree high.

For species diversity in the canopy and offspring, we estimated the Shannon index, *H’*, (Shannon 1948), which considers both species richness and evenness. It was calculated by counting the number of different species and their frequency of appearance (Figure 1), and then applying the following formula (1) where S = species richness and pi = relative abundance of species i.

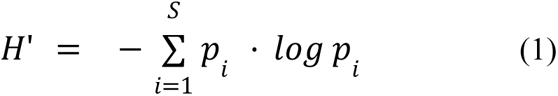

### 2. Genetic analyses

Two genotyping approaches (Figure 2) were used to estimate the genetic diversity of the study populations as well as compare adults and offspring: 1) Sequence-based genotyping using GBS and 2) PCR-based assay for SNP genotyping using Standard Biotools microfluidics SNP genotyping platform (Standard Biotools, San Francisco, CA, USA). The GBS was used for sequence-based genotyping (SNP Set1) and for the SNP discovery needed to perform the PCR-based SNP genotyping (SNP set2). We compared the results from both approaches to validate the use of a small SNP set (SNP set2).

**Figure 2.**
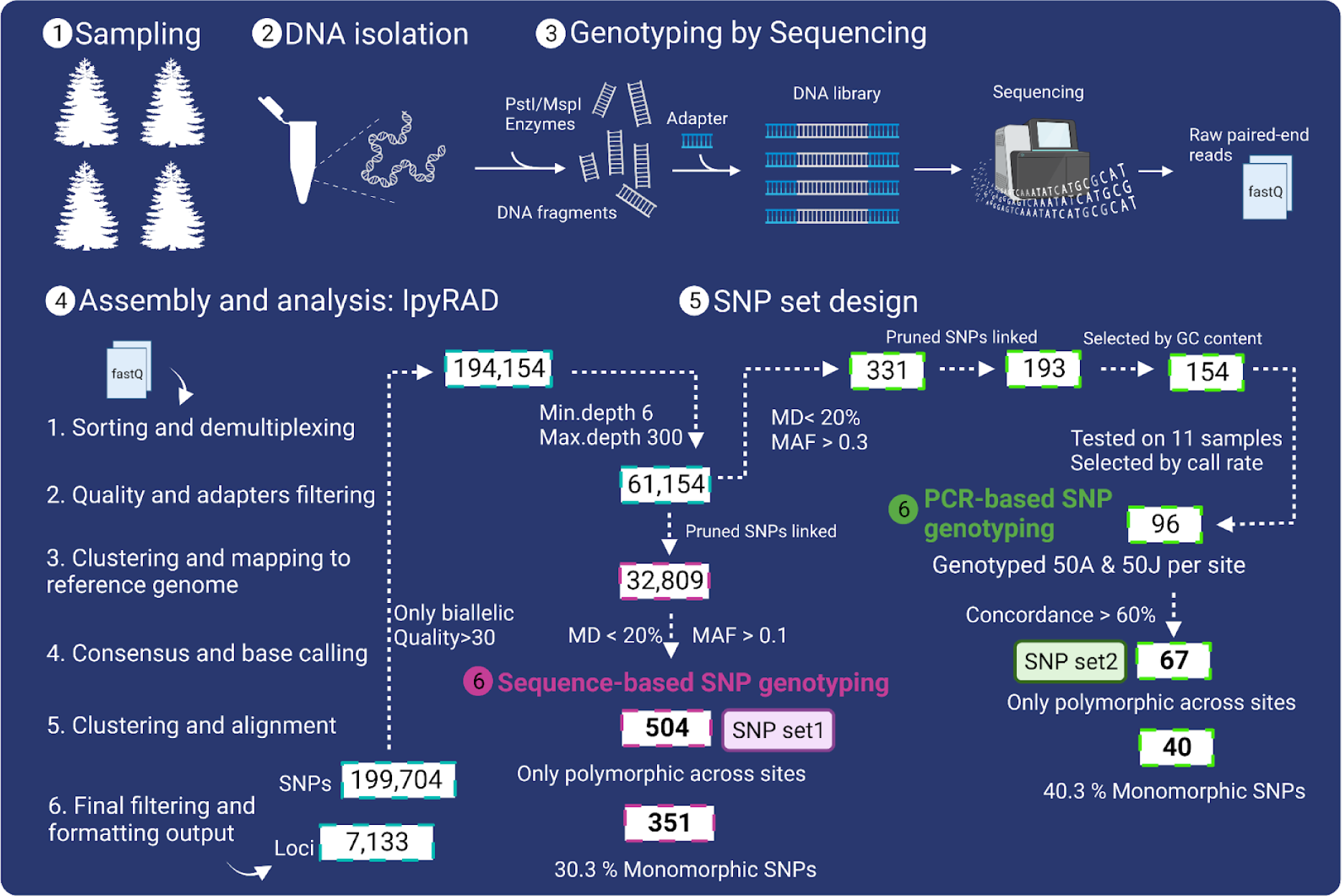
Genetic analysis workflow. An overview is given of each step performed from the sampling (1) and DNA isolation (2), the GBS (3), the assembly and variant calling using IpyRad (4) to the design of both SNP sets (5) to use in the genotyping steps (6, pink: SNP set1, green: SNP set2). MD = missing data, MAF = minor allele frequency.

#### 2.1 Sampling and DNA isolation

We sampled foliage from 34 to 48 adults and 41 to 46 juveniles across sites (Table 5) (Figure 2). To distribute the sampled trees evenly throughout the site area, we calculated random points, resulting in a uniform distribution of both adults and juveniles across the site. The tissue samples were stored in paper envelopes and placed in plastic zip bags containing silica gel beads to promote their desiccation. A subsample of 10 adults and 10 juveniles per site, 80 samples in total, was used to perform the GBS step.

The DNA of the 80 used for GBS was isolated using the spin-column DNeasy plant mini kit (Qiagen USA, Valencia, CA) (Qiagen). For the other 259 samples, we used the temperature-driven enzymatic cocktail Plant DNA Extraction (MicroGEM International PLC 2019) for higher throughput (Guillardín and MacKay 2023). After recovering the DNA we performed a clean-up step by using 0.8X concentration of AMPpureXP beads (Beckman Coulter™, Brea, CA, USA).

The DNA concentration was determined by using Qubit 4 Fluorometer (Invitrogen, Thermo Fisher Scientific, Cleveland, OH, USA). We used NanoDrop One Spectrophotometer (Thermo Fisher Scientific, Wilmington, DE) to estimate the purity based on absorbance ratios A260/A280 and A260/A230. A minimum DNA concentration of 20 ng/ul and absorbance ratios of >1.8 were required for genotyping steps.

#### 2.2 GBS library construction, sequencing and quality check

In GBS, genomic DNA is fragmented by one or more restriction digestion enzymes and then sequenced to uncover polymorphic sites across the genome. To select enzymes producing a suitable cut rate in the library preparation, we analysed the *T. plicata* genome sequence (Shalev et al. 2022) with the simulation software ddRADseqtools (Mora-Márquez et al. 2017). We compared the number and length of fragments recovered from each pair of available enzymes tested (Online resource 1) and selected the optimal combination.

The DNA samples were normalised to 20 ng/ul for library preparation (Genomic Analysis Platform of Laval University, Canada) and sequencing (Genome Quebec, Canada). The library was constructed with PstI/MspI enzyme combination (NEB, Ipswich, MA, USA) and the Illumina TruSeq HT Library Preparation Kit (Illumina, Inc. San Diego, CA, USA) (Figure 2), retaining on the fragments above 300bp from a fragment selection step. Sequencing was performed on an Illumina NovaSeq 6000 PE150 platform (Illumina Inc. San Diego, CA, USA), and 150 bp paired-end raw files were obtained. The fastq files were checked for quality using the FastQC software (S. Andrews 2010); the initial demultiplexing step used SABRE (Barry et al. 1989) based on the barcodes supplied by the Illumina sequencer provider, not allowing any mismatches or Ns, and the Illumina adapters were trimmed using Trimmomatic (Bolger et al. 2014).

#### 2.3 Clustering, assembly and variant calling using Ipyrad

We used the Ipyrad pipeline (Eaton and Overcast 2020) to cluster, assemble and call the bases from the trimmed fastq files (Online resource 2). Ipyrad is divided into six steps (Figure 2), allowing the repetition of steps separately once the results have been reviewed. After the first Ipyrad run including all the samples, we performed exploratory depth, missing data and number of loci per sample tests. We excluded the samples that showed an average depth below 6 reads or more than 50% of missing data. We reran Ipyrad from the 3rd step without the excluded samples. All the runs were performed in the University of Oxford Advanced Research Computing (Oxford, UK) facility and used a Capability cluster node with 48 CPUs and 384GB RAM.

#### 2.4 Data post-filtering and SNP sets design

A subset of informative SNP markers was developed for the general use in research and conservation in *T. plicata* with a two-step filtering process described in Figure 2.

First, pre-filtering of the initial GBS dataset was done to remove multi-allelic SNPs (--max-alleles 2) using PLINK (Purcell et al. 2007) and indels (--remove-indels) followed by a read-depth pre-filtering per locus between 6 and 300 (--min-meanDP --max-meanDP) and a minimum quality of 30 across the SNPs (--min quality 30) using VCFtools software (Danecek et al. 2011). Second, a post-filtering procedure was used to identify a set of SNPs distributed across the genome, which were polymorphic and had low missing data. We followed more stringent filtering criteria to design the reduced PCR-based SNP genotyping set compared to the sequence-based SNP genotyping to obtain the desired number of 96 SNPs to test in the genotyping platform (Figure 2).

##### Sequence-based genotyping (SNP set1)

The GBS SNP genotyping used standard filtering criteria to reduce the missing data while ensuring that the markers are distributed across the genome and have sufficient polymorphism within and among sites. To develop a sequenced-based genotyping SNP set (SNP set1), we pruned the SNPs under strong linkage disequilibrium by using PLINK (Purcell et al. 2007), calculating the pairwise correlation in a 50-SNP window and discarding SNPs with a correlation higher than 0.1, in increments of 10 SNPs at a time (-indep pairwise 50 0.1 10) (Figure 2).

Next, we uploaded the resulting VCF file into RStudio (R Core Team 2018; RStudio Team 2020) using the read.vcfR function from the vcfR package (Knaus and Grünwald 2017) to convert it into a genlight object for processing in the dartR (Mijangos et al. 2022) package environment using the vcfR2genlight tool. DartR facilitates the import and analysis of SNP data and includes the use of 3rd party population genetics software. Using the gl.filter.callrate function, we kept only the individuals with less than 50% of missing data and the SNPs that showed less than 20% of missing data. To ensure that the genetic variants are sufficiently common and informative, we used a cut-off of minor allele frequency (MAF) <0.1 with gl.filter.maf function across the whole dataset. We quantified the monomorphic SNPs on each site using gl.filter.monomorphs.

##### PCR-based genotyping (SNP set 2)

The PCR-based SNP set (SNP set2) followed a similar approach but other filters were applied. After the read-depth pre-filtering, we used VCFtools (Danecek et al. 2011) to filter out SNPs with missing data above 20 % (--max-missing) and a MAF < 0.3 (--maf). Then, we pruned strongly linked SNPs using PLINK (Purcell et al. 2007) with the same criteria as for the SNP set 1. We selected 100 bases on each side of the variant to retrieve the SNP sequences by using *T. plicata* reference genome (v3) (Shalev et al. 2022) in the Python package PySAM (Gilman et al. 2019). The sequences thus selected were used to design the assays and perform a quality assay check by Standard Biotools (San Francisco, CA, USA). The assays were filtered to discard sequences with GC>65%. The resulting SNP-type assays were tested on 11 samples by using FlexSix IFC plates on the Standard Biotools microfluidics SNP genotyping platform using the Juno system to load and combine the microfluidics, the Biomark to perform the PCR, and the FluidiGM SNP Genotyping software to digitalised and analyse the image outputs (all from Standard Biotools, San Francisco, CA, USA). We used Fast Probe Master Mix (Biotium Inc. Fremont, CA) and followed the recommended preparation protocol (Standard Biotools 2015b). We selected the best 96 SNP assays based on the level of polymorphisms and call rate to genotype the total sample of 339 individuals. We used eight 192.24 IFC plates following the manufacturer protocol (Standard Biotools 2015a). The results were visualised using the FluidiGM SNP genotyping software and exported to an SNP table file.

We ran a consistency test against the SNP set1 comparing the results for the 80 samples and the SNPs in common, following a modified estimation error method using duplicate samples as described in Mastretta-Yanes et al. 2015. We discarded SNPs with a concordance < 60%. Finally, we quantified the number of exclusive monomorphic SNPs as these are expected to indicate large differences in some SNPs MAF among sites.

#### 2.5 SNP set validation

The SNP set2 markers were validated to ensure their accuracy and reliability for subsequent genetic diversity analyses by comparing results obtained in population structure analyses

From both sets, we excluded the private monomorphic SNPs of each site to avoid bias in genetic structure due to site-exclusive monomorphic SNPs (Nazareno et al. 2017; Schmidt et al. 2021). We assessed the ability of both SNP sets to identify population structure using the Discriminant analysis of principal components (DAPC) (Jombart et al. 2010) and the Principal Coordinate Analysis (PCoA) methods (Zuur 2007). We used the dapc function from adegenet R package (Jombart 2008) and the gl.pcoa function to implement the analysis and gl.pcoa.plot from dartR (Mijangos et al. 2022) to visualise the results. To compare the level of genetic differentiation among sites, we used Arlequin (Excoffier and Lischer 2010) to estimate the sites’ F_st_ pairwise genetic distances, and we performed hierarchical Analyses of Molecular Variance (AMOVA) (Meirmans 2012) to evaluate the amount of population genetic variation within and among sites.

#### 2.6 Population genetics analysis

##### Genetic diversity and genetic variation

We estimated the genetic diversity within each site and compared adults and juveniles with data from SNP set2. Estimates were made for two scenarios, including and excluding private monomorphic SNPs following the recommendations of Schmidt et al. (2021). We estimated observed heterozygosity (H_o_) and expected heterozygosity (H_e_) using the gl.report.heterozygosity tool in DartR package (Mijangos et al. 2022). Significant differences among expected heterozygosity between adults and juveniles within sites were tested using the tool gl.test.heterozygosity. We used the tool divBasic from the package diveRsity (Keenan et al. 2013) to calculate the inbreeding coefficient (F_is_) executing 1000 bootstraps to estimate the 95% confidence intervals. We compared the contemporary effective population size (N_e_) of adults and juveniles using only monomorphic SNPs (40 SNPs in Longleat1 and 67 SNPs in the other sites) using the linkage disequilibrium method implemented in NeEstimator V2.1 software (Do et al. 2014) and used the Jackknife method to estimate the confidence intervals (Jones et al. 2016).

##### Kinship analysis within sites

To assign offspring to putative parents we used both monomorphic and polymorphic SNPswith the software CERVUS (Kalinowski et al. 2007). We removed SNPs not in Hardy-Wineberg equilibrium (HWE) in each site to focus on relationships that are less influenced by biological processes like inbreeding or selection, which can lead to deviations from HWE. The final number of SNPs used in every site was over 60 (Table 6), which has been suggested as the minimum number of SNPs to accurately estimate kinship relations (Anderson and Garza 2006). We used the GPS waypoints of each individual to determine parent-offspring distances.

## Results

### 1. Study sites characterisation: structure and species diversity

The study sites were characterised in terms of area, isolation status, stand distribution, canopy cover, tree density, *T. plicata* specific tree density, and average height (Table 1). The total tree density ranged from 206.9 to 373 trees/ha; Stourhead2 had the largest difference between the overall tree density (226.4 trees/ha) and the *T. plicata* density (21.2 trees/ha) (Table 1). We found variation across sites in regard to size, with Stourhead2 being the largest, patchiness due to trees being in smaller compartments within the sites that were planted in the same year (information provided by owner), canopy cover and average tree height (Table 1).

Stourhead2 had a high canopy species diversity (*H’* = 1.5), while the other three sites were all below 0.8 (Figure 3 panel A). The offspring species diversity ranged from 0.9 in Bagley Wood to 1.8 in Stourhead2. The distribution of tree diameter classes showed that Stouhead2 had a reverse-j profile, characteristic of woodlands in a late stage of transformation to CCF, and only one peak was found in the other sites (Figure 3 panel B and C). The structural diversity and levels of transformation (Figure 3 panel B) was consistent with the levels of species diversity (Figure 1 panel A).

**Figure 3.**
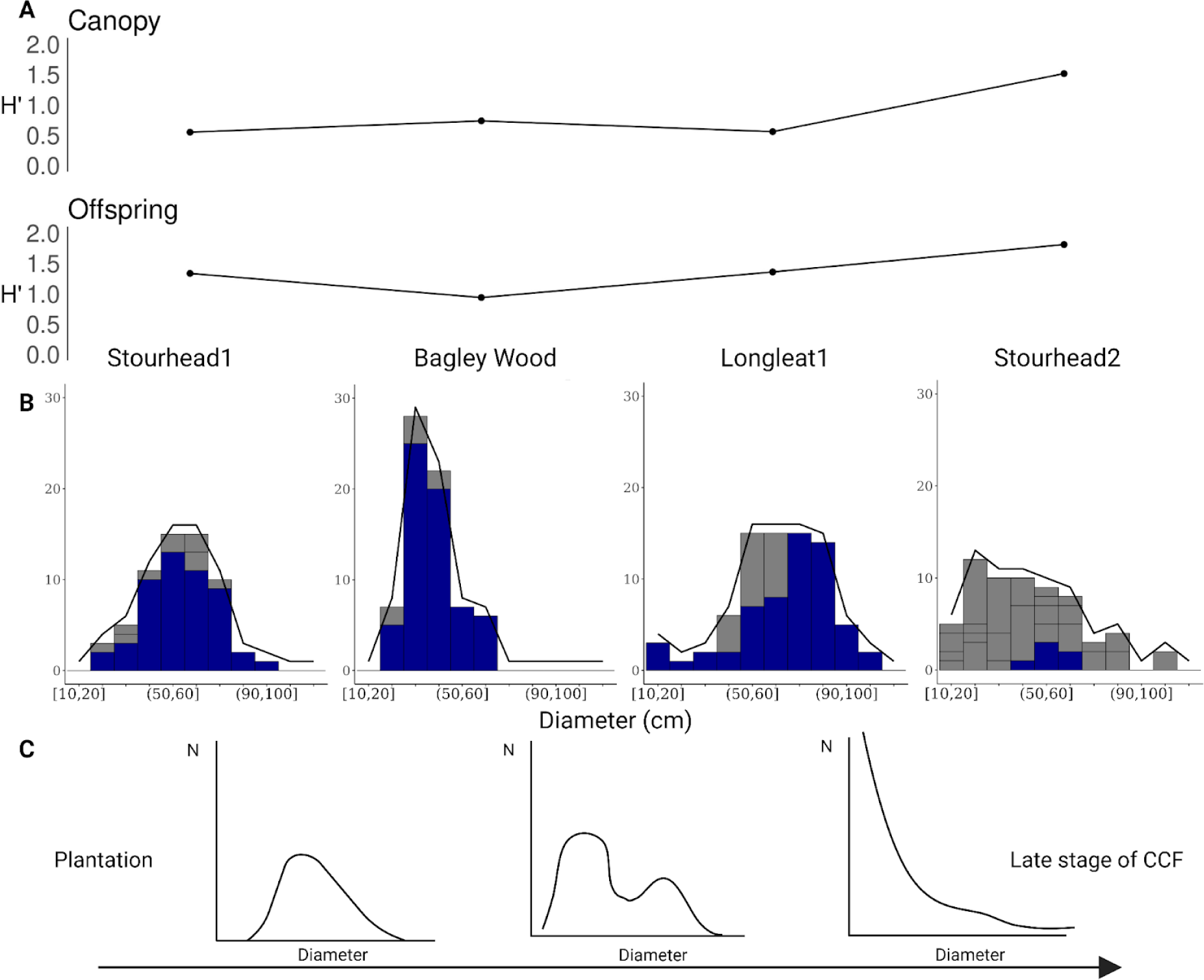
Structural and Species diversity. A) Shannon index (*H’*) of the canopy and offspring per site, B) Structural diversity per site shown as the number of trees per diameter class, *T. plicata* individuals coloured in blue, C) Structural diversity transformation from plantation to CCF late stage, modified from Pommerening and Murphy, (2004).

### 2. DNA isolation, sequencing, quality check and SNP calling

Simulations using ddRADseqTools (Mora-Márquez et al. 2017) and the reference genome (Shalev et al. 2022) were performed to predict which enzymes would yield a number of DNA fragments to ensure high genome coverage in the GBS analysis (Online resource 1). The restriction enzymes PstI/MspI gave 112,191 fragments and were selected to construct the library (see methods for library construction and sequencing details). The enzymes ApeKI, BfaI and NsIi/MspI were estimated to produce >300,000 fragments, and SbfI/MspI was estimated at <10,000 fragments of 125-300bp size, which appeared as too high and too low coverage of the *T. plicata* genome, respectively (Online resource 1). Total DNA isolated from 80 individuals was used for GBS analysis (see methods for details). The FastQC analysis of the resulting raw reads gave high-quality scores with an average Phred quality score above 30. We trimmed the 3’ end of the sequences and removed the adapter sequences.

After a first run of the assembler and base calling with the software Ipyrad (Eaton and Overcast 2020), three samples had read depth <3.5 and two other samples had >70% of missing data. The remaining 75 samples were retained for further data analysis, which included 1,273,904,871 raw reads (Online resource 3), and identified 199,704 SNPs variants called in 7,133 loci.

### 3. Data post-filtering and SNP set design

We recovered 61,154 SNPs within 4,134 loci with an average read-depth of 100.5 across loci (range of 6-300), after filtering to remove indels, multi-allelic SNPs and low-quality SNPs (see methods for details).

The sequenced-based SNP set was then reduced to 504 SNPs (SNP set1) by filtering for missing data and distribution of the SNPs across the genome and the level of polymorphism. Starting from the initial 61,154 SNPs, strongly linked SNPs were pruned, and samples with >50% of missing data and SNPs with >20% of missing data were removed, resulting in a dataset of 66 samples and 9,343 SNPs in 1,983 loci. A final filtering for MAF >0.1 across the four sites reduced the set to 504 SNPs across 443 loci (Table 2). We detected 153 private monomorphic SNPs (30.3%) (Table 2) and explored their distribution by site showing an unbalanced distribution, with above 76% of the monomorphic SNPs only in Longleat1 (Figure 4).

**Figure 4.**
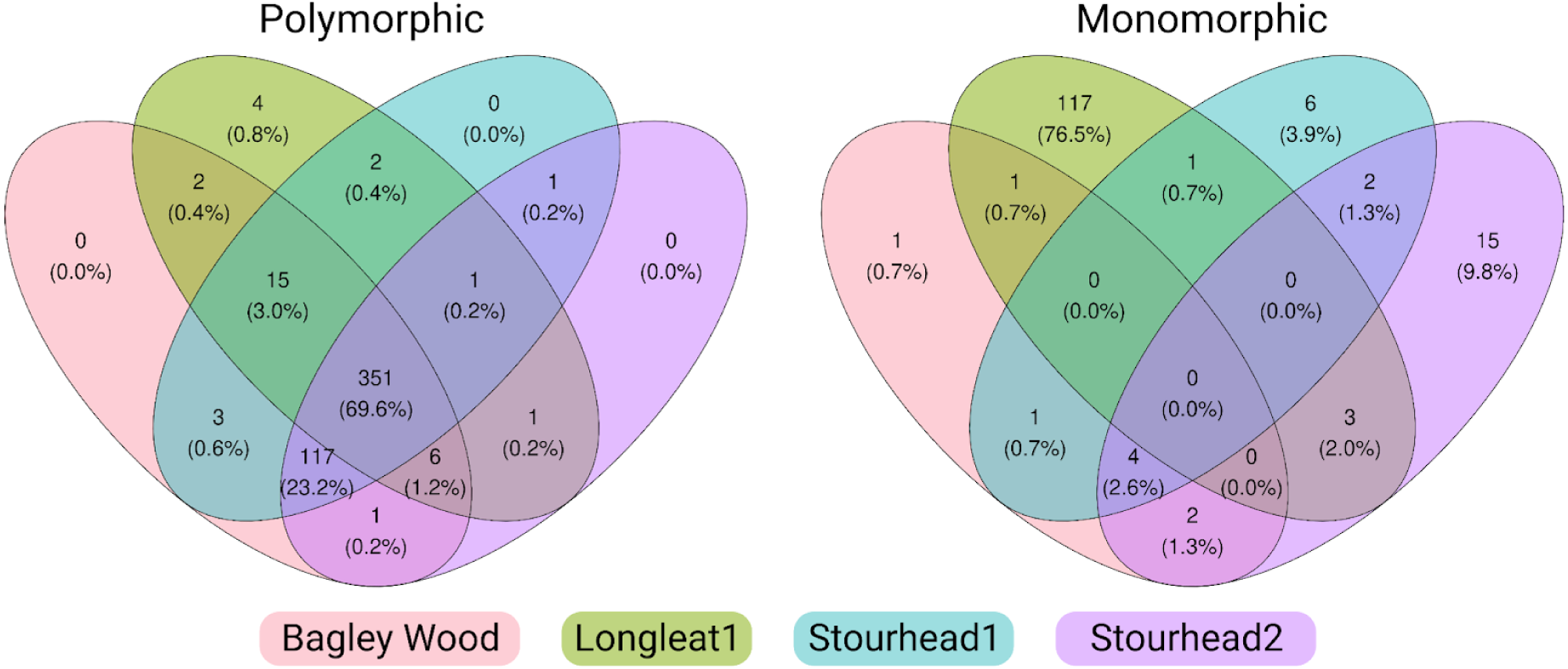
SNP set1 private polymorphic and monomorphic SNPs distribution across sites.

**Table 2.** Description of the two SNP sets developed in this project. N: Number.

|  | N SNPs | N private Monomorphic SNPs | N samples genotyped | Mean missing data % | Mean minor allele frequency |
| --- | --- | --- | --- | --- | --- |
| SNP set1 | 504 | 153 | 75 | 8.53 | 0.280 |
| SNP set2 | 67 | 27 | 360 | 4.23 | 0.397 |

We developed a PCR-based genotyping assay from the output of the GBS analysis and obtained data from 67 SNPs (SNP set2) across the full sample set, with pre-filtering and post-processing selection steps. We identified 193 SNPs by filtering as follows, <20% missing data and those that had MAF>0.3 and pruned the SNPs strongly linked, resulting in 193 SNPs in 187 loci. We further excluded 39 SNPs based on their flanking sequences due to GC >65% (12 SNPs) or high levels of repetitive regions (27 SNPs). The remaining 154 pairs of primers were tested using the Standard Biotools microfluidics SNP genotyping platform (Standard Biotools, San Francisco, CA, USA). We selected the best 96 assays based on call rate and polymorphism criteria to genotype all 339 samples. A consistency test was performed on samples tested in duplicate and 25 SNPs with <60%, resulting in a final set of 67 SNPs (SNP set2). We found that 27 SNPs (40.3%) (Table 2) were private monomorphic to Longleat1 and were exclusively found in the adults.

### 4. SNP set validation

We assessed the ability of both SNP sets 1 and 2 to capture the genetic structure among sites using both a DAPC and a PCoA method and found patterns of population structure (Figure 5, Online resource 4). Axis 1 of the DAPC (89.8% and 90.1%) of both tests showed a genetic differentiation of Longleat1 against the other three sites; PCA axis1 (12.3% and 9.2%) followed the same pattern, although the variation explained by the test is lower.

**Figure 5.**
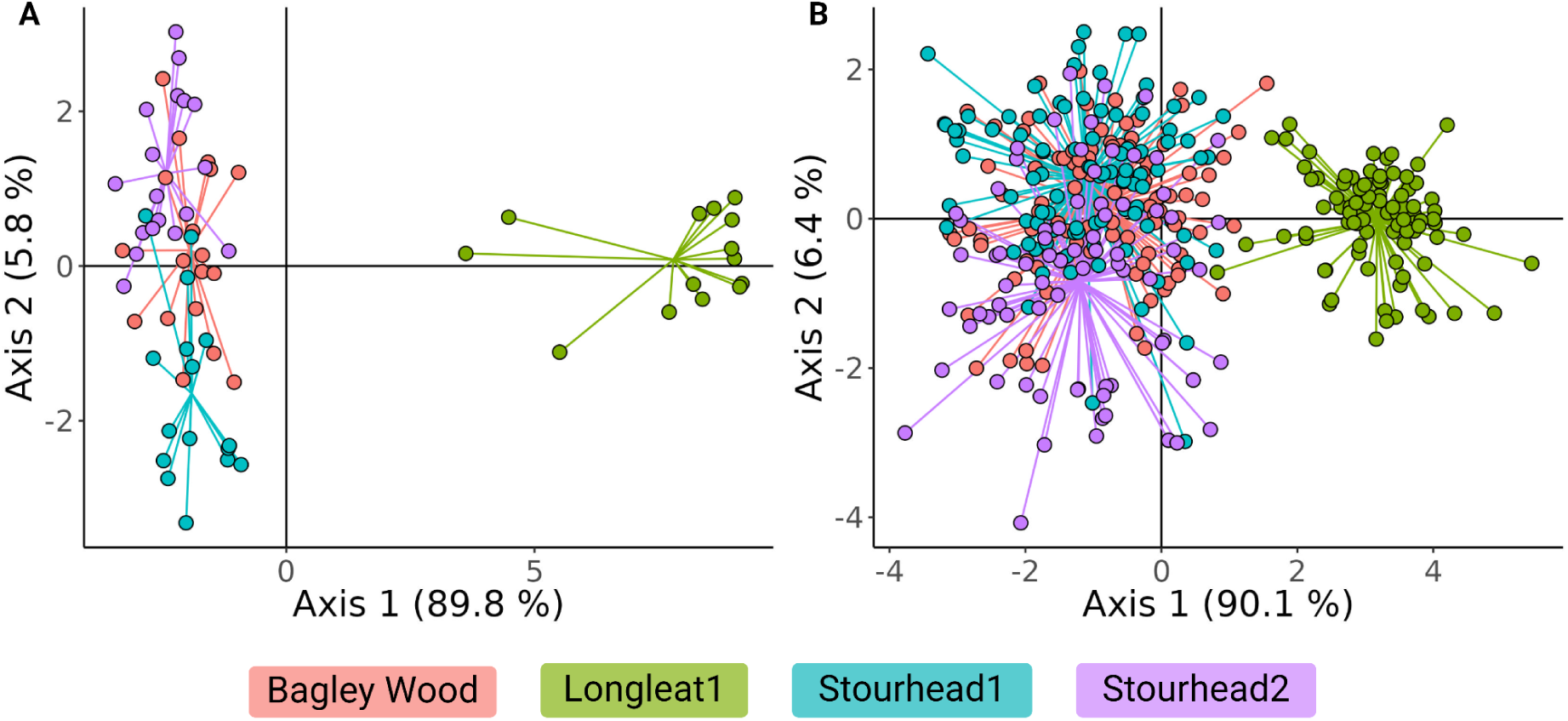
Discriminant Analysis of principal components (DAPC) results using A) SNP set1 and B) SNP set2 across the four study sites.

Analyses of molecular variance (AMOVA) were used for partitioning of genetic variation among and within the study sites (Table 3). For SNP set1, the analysis revealed that 6.98% of the total genetic variation is attributed to differences among the four sites while SNP set2 showed 11.34%, both with a significant p-value (<0.001) indicating genetic differentiation among the populations. The remaining genetic variation is found within the sites (93.02% and 88.66%) also with significant p-value (<0.001) (Table 3).

**Table 3.** Comparison of results from SNP set 1 vs SNP set2 components of molecular variation (AMOVA) for the four study sites. Significant P-value <0.05

|  | Source of variation | d.f | Sum of Squares | Percentage of variation | Pvalue |
| --- | --- | --- | --- | --- | --- |
| SNP set1 | Among sites | 3 | 274.886 | 6.98 | $<0.01$ |
| | within sites | 716 | 4535.281 | 93.02 | $<0.01$ |
|  | Total | 719 | 4810.167 |  |  |
| SNP set2 | Among sites | 3 | 366.065 | 11.34 | <0.01 |
|  | within sites | 128 | 3000.049 | 88.66 | <0.01 |
|  | Total | 131 | 3366.114 |  |  |

In the pairwise F_st_ comparisons between sites (Table 4), Longleat1 had the highest genetic differentiation with a pairwise F_st_ >0.1 when compared to every other site, with both SNP sets (p-values <0.001). The lowest pairwise F_st_ was found between Bagley Wood and Stourhead1, F_st_ = 0.01211 and F_st_ = 0.00234 with SNP set1 and SNP set2, respectively, which are non-statistically different. Taken together, the data show very good concordance between the SNP sets.

**Table 4.**
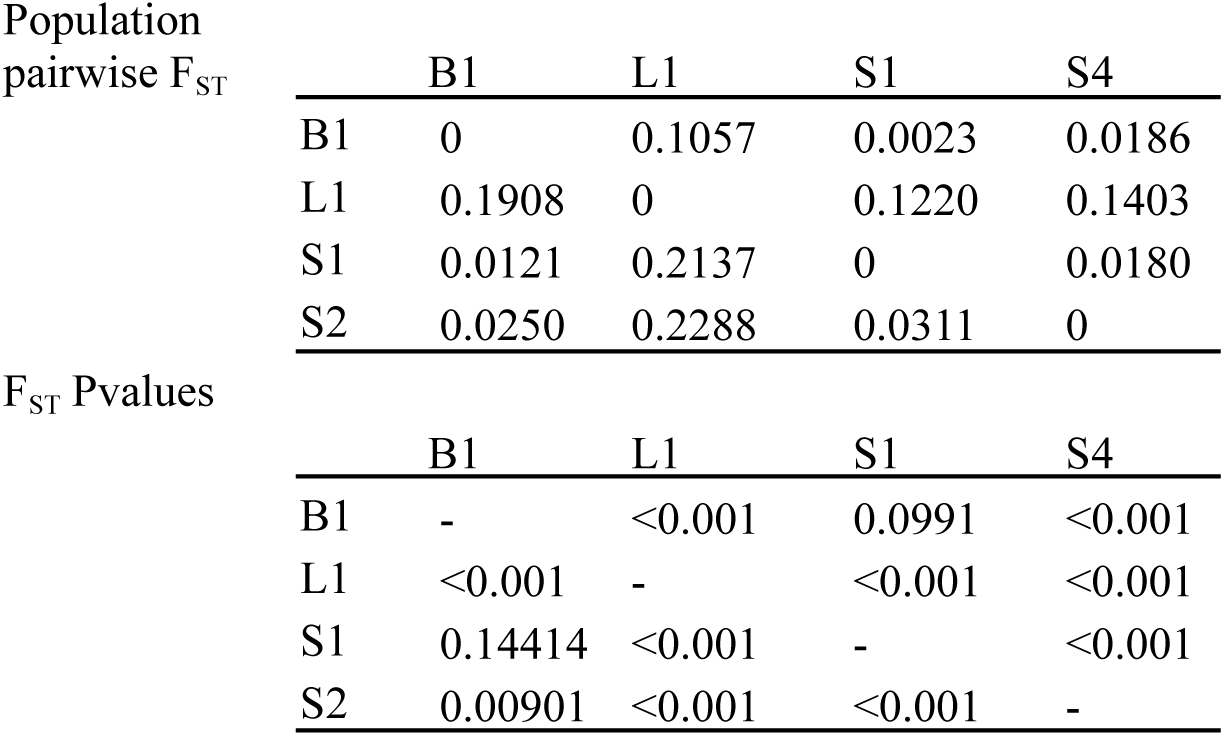
F_ST_ pairwise site comparisons. Above the diagonal, results of SNP set2, below diagonal, results of SNP set1. d.f = degree of freedom. Significant P-value <0.05.

### 5. Population genetic analyses

#### 5.1 Genetic diversity and genetic variation

The genetic diversity indicators (Table 5) compare the results when using only the polymorphic SNPs to results when using both the monomorphic and polymorphic SNPs. Overall, we found high heterozygosity levels across and within populations. The observed heterozygosity (H_o_) ranged from 0.255 to 0.460 across sites when using all makers, with the highest observed in Bagley Wood juveniles and the lowest in Longleat1 juveniles. With the polymorphic SNPs alone, H_o_ generally increased across sites, with the highest H_o_ in Longleat1 adults (0.408) and the lowest in Stourhead2 juveniles (0.338). The largest change when excluding the monomorphic SNPs was observed in Longleat1 adults and juveniles (H_o_ from 0.261 to 0.437 and 0.255 to 0.386, respectively). We found significant differences in expected heterozygosity (H_e_) between adults and juveniles in Longleat1 (0.259 and 0.291) and Stourhead2 (0.439 and 0.456) using both monomorphic and polymorphic SNPs; however, the difference only held for Stourhead2 adults and juveniles (0.441 and 0.462) when the monomorphic markers were excluded.

**Table 5.** Genetic diversity results. Significant differences were tested between adults (A) and juveniles (J) within each site. Significant p-value <0.05 is shown in the table as “*”. N_s_ = number of samples collected, C.I = confidence intervals, S.D = standard deviation, H_o_ = observed heterozygosity, H_e_ = expected heterozygosity, F_is_ = Inbreeding index.

|  | Bagley Wood |  | Longleat1 |  | Stourhead1 |  | Stourhead2 |  |
| --- | --- | --- | --- | --- | --- | --- | --- | --- |
|  | A | J | A | J | A | J | A | J |
| $N_s$ | 48 | 44 | 44 | 45 | 37 | 46 | 34 | 41 |
| Including monomorphic |  |  |  |  |  |  |  |  |
| $H_o$ | 0.398 | 0.393 | 0.261 | 0.255 | 0.367 | 0.410 | 0.381 | 0.337 |
| $H_o$ S.D | 0.084 | 0.086 | 0.249 | 0.192 | 0.096 | 0.099 | 0.097 | 0.088 |
| $H_e$ | 0.460 | 0.459 | 0.259 | 0.291* | 0.448 | 0.453 | 0.439 | 0.456* |
| $H_e$ S.D | 0.041 | 0.041 | 0.243 | 0.210 | 0.056 | 0.058 | 0.070 | 0.060 |
| $F_{is}$ | 0.134 | 0.144 | -0.007 | 0.125 | 0.179 | 0.094 | 0.132 | 0.262 |
| lower C.I | 0.086 | 0.078 | -0.087 | 0.049 | 0.095 | 0.048 | 0.073 | 0.162 |
| upper C.I | 0.182 | 0.217 | 0.076 | 0.194 | 0.263 | 0.142 | 0.191 | 0.350 |
| Only polymorphic |  |  |  |  |  |  |  |  |
| $H_o$ | 0.407 | 0.408 | 0.437 | 0.386 | 0.372 | 0.419 | 0.377 | 0.338 |
| $H_o$ S.D | 0.081 | 0.089 | 0.162 | 0.132 | 0.106 | 0.092 | 0.095 | 0.076 |
| $H_e$ | 0.463 | 0.461 | 0.434 | 0.440 | 0.451 | 0.455 | 0.441 | 0.462* |
| $H_e$ S.D | 0.041 | 0.039 | 0.149 | 0.132 | 0.057 | 0.055 | 0.070 | 0.049 |
| $F_{is}$ | 0.122 | 0.115 | -0.007 | 0.124 | 0.174 | 0.081 | 0.144 | 0.268 |
| lower C.I | 0.071 | 0.044 | -0.088 | 0.049 | 0.083 | 0.026 | 0.060 | 0.178 |
| upper C.I | 0.171 | 0.188 | 0.074 | 0.201 | 0.260 | 0.134 | 0.227 | 0.360 |

The inbreeding coefficient (F_is_) was less influenced by the makers used and ranged from -0.007 to 0.262 for both monomorphic and polymorphic SNPs and - 0.007 to 0.268 for only polymorphic (Table 5). Longleat1 had the lowest inbreeding coefficient and Stourhead2 had the highest, and no significant differences were found between adults and juveniles within sites. The levels of N_e_ ranged from very low (N_e_ < 10) in Stourhead2 (adults and juveniles) and Longleat1 (adults) to low (N_e_ < 100) in all the other sites (Table 6). Juveniles in Stourhead1 and Longleat1 were noted for having substantially higher N_e_ compared to adults.

**Table 6.**
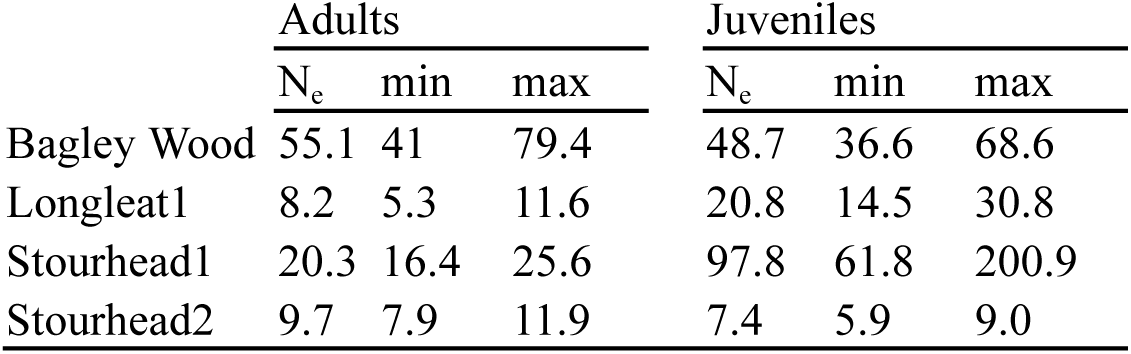
Effective population size (N_e_) estimated in adults and juveniles across sites. min and max correspond to the 95% confidence intervals calculated using the Jackknife method (Jones et al. 2016).

|  | Adults |  |  | Juveniles |  |  |
| --- | --- | --- | --- | --- | --- | --- |
|  | N<sub>e</sub> | min | max | N<sub>e</sub> | min | max |
| Bagley Wood | 55.1 | 41 | 79.4 | 48.7 | 36.6 | 68.6 |
| Longleat1 | 8.2 | 5.3 | 11.6 | 20.8 | 14.5 | 30.8 |
| Stourhead1 | 20.3 | 16.4 | 25.6 | 97.8 | 61.8 | 200.9 |
| Stourhead2 | 9.7 | 7.9 | 11.9 | 7.4 | 5.9 | 9.0 |

#### 5.2 Kinship analysis

The assignment of individuals in the natural regeneration to their parents ranged from 11% in Longleat1 to 37% in Bagley Wood, and showed trends consistent with individual site properties. The average distance between assigned parents and offspring across sites is 192.28 m. Stourhead1 is a patchy stand (Table 1) and had both the lowest average distance (65.8 m) and the lowest maximum distance (310.4 m) across sites (Table 7). Stourhead2 had the lowest *T. plicata* density (21.2 trees/ha) and the largest surface area (18.57 ha) (Table 1); it also had the largest minimum assignment distance across all sites (15.9 m). Bagley Wood is a patchy stand (Table 1); it had the largest average and maximum distance of assigned parents-offspring and the most parents-offspring assigned (Table 7). Longleat1 had the lowest number of parents-offspring assigned (11%) and the parent assigned to the highest number of offspring (Table 7).

**Table 7.**
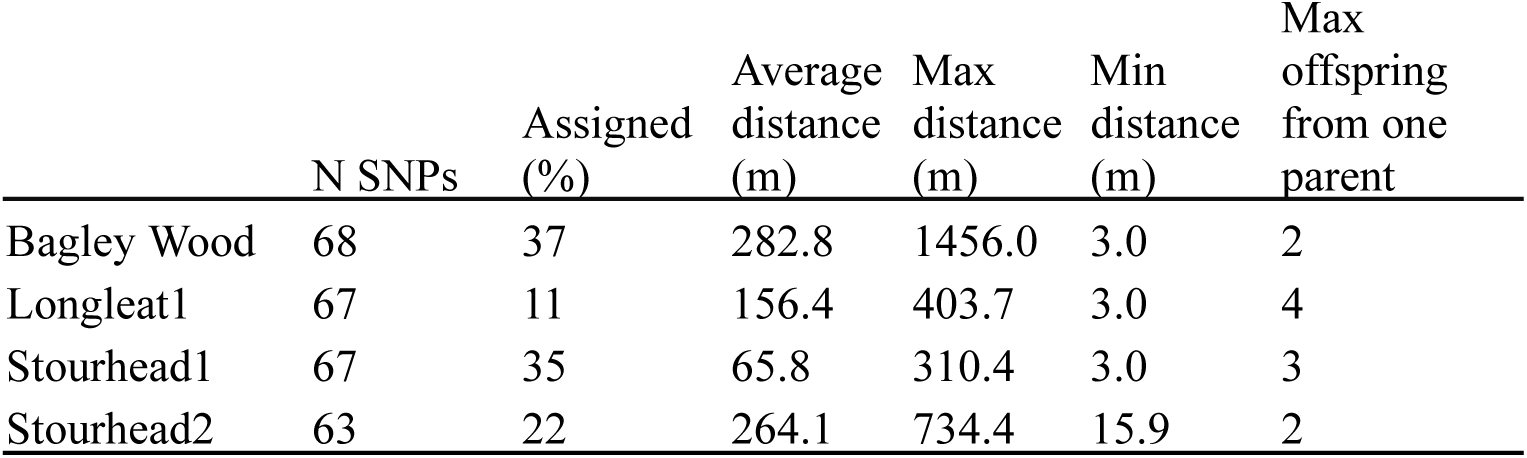
Kinship analysis results, including the number of SNPs used per site (N SNPs), the percentage of parent-offspring related found (Average %), the mean, maximum and minimum distance these relationships were found and the maximum number of offspring related to one parent found (Max offspring from one parent). N: number, Max: maximum, Min:minimum.

## Discussion

In this study we assessed the genetic diversity in planted *T. plicata* trees and their offspring in four UK woodlands managed under the CCF approach. We identified 504 informative SNP markers (SNP Set1) by using a GBS approach and designed a reduced SNP set (67 SNPs) (SNP Set2) for genotyping on a PCR-based platform. We found significant differences in genetic variation and in the number of private monomorphic SNPs among the study sites. Overall, the heterozygosity levels were high (H_e_ and H_o_) across and within sites. Only one site (Stourhead2) showed significantly higher H_e_ in the juvenile trees compared to adults. However, we found very low N_e_ across and within sites, with adults ranging from 8.2 in Longleat1 to 55 in Bagley Wood, and juveniles ranging from 7.4 in Stourhead2 to 97.8 in Stourhead1, which may indicate low levels of genetic diversity and potential adaptability.

### 1. Sites biodiversity and level of transformation

The four woodlands planted with *T. plicata* varied in tree density, canopy coverage, structure and species diversity (Table 1). To define the stands’ CCF level of transformation, we used traditional forestry survey methods estimating species and structural diversity; however, new high-throughput methods using unmanned aerial vehicles (UAV) are becoming popular for this type of data collection (Bennett et al. 2020).

Stand thinning practices such as opening gaps are often used to initiate the transformation toward CCF to encourage natural regeneration (Kerr et al. 2017). Increased species diversity in the stand indicates the development from pure even-aged plantation into continuous cover forest woodlands (Gärtner and Reif 2004). We found a trend in our sites where higher number of species was linked to more developed structural diversity, and therefore, we were able to assign different levels of CCF transformation as described in several studies (Bravo-Oviedo et al. 2014; Gärtner and Reif 2004; Kerr et al. 2010, 2017). The canopy species diversity as indicated by the Shannon index ranged from 0.54 in Bagley Wood to 1.5 in Stourhead2, compared to other UK stands in transition into CCF studied by Pommerening, (2002), which ranged from 0.01 to 0.62.

Changes in tree diameter distribution from even-aged plantation to late stage of CCF were described in Pommerening and Murphy, (2004) and Vitkova and Dhubháin, (2013). Bagley Wood, Longleat1 and Stourhead1 had a structure similar to pure plantations composed of > 90% studied in *Pinus sylvestris* and *Picea abies* by Zaytsev et al. (2019). Alternatively, Stourhead2 had a reverse-j structure typically found in unmanaged or near-natural woodlands (De Quesada and Kuuluvainen 2020; Kerr 2002) and late-stage CCF forests (Kerr et al. 2010, 2017). We found a low overall tree density ranging from 206.9 in Longleat1 to 373.0 in Bagley Wood compared to sites studied by Kerr et al. (2017) which ranged from 557 to 716 trees/ha. The sites in Kerr et al. (2017) have different tree species including *T. plicata* and have been managed using the Bradford-Hutt system, which is used to transform even-aged plantations into CCF focusing on managing small areas actively using planting, thinning and felling practices.

Overall, Stourhead2 gave the highest biodiversity and the most diverse structure showing a heterogeneous mix of species and sizes. The sites at Bagley Wood, Longleat1 and Stourhead1 shared similar levels of transformation into CCF and similar canopy species diversity. Bagley Wood had the lowest seedling diversity, which may compromise the future canopy diversity on this stand. Further monitoring of the effects of low species and structure diversity may contribute to understanding the success factors when transforming pure even-aged plantations into CCF woodlands.

### 2. Low number of variants and monomorphic SNPs

Genotyping by sequencing can discover thousands of genomic variants in non-model species but requires a bioinformatic procedures for data cleaning due to genotyping errors (Mastretta-Yanes et al. 2015) and missing data (López de Heredia et al. 2020; Vaux et al. 2023). This typically includes steps to filter or test for missing data, minor allele frequency, minimum and maximum depth, hardy-weinberg and linkage disequilibrium filtering (Vaux et al. 2023), and may involve imputation of missing alleles using statistical methods (López de Heredia et al. 2020; Mora-Márquez et al. 2023). We found 199,704 variants before filtering, which is relatively low compared to the 1.3M SNPs found in Chinese fir (*Cunninghamia lanceolata*) also a member of the Cupressaceae family (Zheng et al. 2019) or the 349,542 SNPs found in the foxtail pine (*Pinus balfouriana*) (Friedline et al. 2015). We selected a set of high-quality SNPs to design a reduced marker set with stringent filtering criteria (see methods section). After filtering by depth, MAF, missing data and pruning the loci linked, we retained 504 high-quality SNPs which exhibited a high proportion of private monomorphic SNPs (30.3%) (Table 2), the majority being found in Longleat1 (Figure 4). The MAF>0.1 filter usually removes rare, private and monomorphic SNPs across the population studied. In our case, 70% (Figure 4) of the site-exclusive monomorphic SNPs are found in Longleat1, indicating a high level of fixed alleles compared to the other sites. Most private alleles were found in the adults revealing low genetic variance and suggesting differences among the seed source provenance used when establishing the plantations.

### 3. SNP genotyping assay development

Our study aim was to retain a reduced set of high-quality SNPs markers selected using stringent filtering criteria (see methods section). The validation of the SNPs discovered using GBS and filtered to perform population genetic analysis followed a similar approach to several other genomic studies (Choi et al. 2020; Kishor et al. 2020; Nguyen et al. 2020). The final 67 SNPs (SNP set2) were validated by testing and comparing the genetic variation and genetic differentiation among sites found using the larger SNP set (SNP set1). The DAPC, PCoA and the AMOVA results using both SNP sets, with only the polymorphic SNPs, showed the same genetic variation, and genetic clustering (Table 3, Figure 5 and Online Resource 4) supported by the F_st_-pairwise results (Table 3). Comparison of results indicated that Longleat1 had the highest F_st_-pairwise values (F_st_ > 0.19) across all sites, and that Stourhead1 and Bagley Wood had no significant genetic differences. This approach is similar to Kishor et al. (2020), who developed a SNP set in melon cultivars and validated their approach by comparing the genetic differentiation in a large data set of 5,644 SNPs and a smaller SNP set (184 SNPs). The SNP set2 may facilitate further genetic studies on *T. plicata* to investigate woodland genetic diversity, and will be particularly useful for large sample sets when GBS may be too costly or very complex to analyse.

### 4. Population genetic analysis

Our data show that heterozygosity levels (both H_o_ and H_e_) were lower across the sites when monomorphic SNPs were included along with the polymorphic SNPs (SNP set 2), most remarkably in Longleat1 (Table 5). This same pattern was found in Schmidt et al. (2021) when they compared the autosomal heterozygosity (including monomorphic SNPs) to the SNP heterozygosity (only polymorphic SNPs) in two species of insects using ddRADseq data. They argued in favour of reporting both results as different biases can affect the genetic diversity estimations when using only polymorphic SNPs, such as the amount of missing data and a small number of samples.

Overall, H_o_ levels across sites were relatively high, ranging from H_o_ = 0.337 in Stourhead2 to H_o_ = 0.410 in Stourhead1, in contrast to the low levels of H_o_ = 0.219 found in the natural range of *T. Plicata* (Shalev et al. 2022). Only Longleat1 showed similar levels than those of Shalev et al. (2022) with H_o_ = 0.259 and H_o_ = 0.291 in adults and juveniles, respectively. The high S.D. found in Longleat1 indicated that the heterozygosity results are dispersed and may suggest variability across loci (Høy Hansen et al. 2022). Low levels of H_o_ in Longleat1 was explained by the large number of monomorphic SNPs found, which could suggest the potential fixation of alleles or high levels of inbreeding in the source populations. Generally, the inbreeding levels remained unchanged after excluding monomorphic SNPs from the analysis and no differences between adults and juveniles were found. However, in three of the sites, the F_is_ in juveniles was higher than in adults, which may indicate a trend that could be of concern for the future. However, the F_is_ ranged from -0.007 in Longleat1 adults to 0.262 in Stourhead2 juveniles, indicating lower inbreeding in these planted woodlands than in the natural range (F_is_ = 0.331), estimated by (Shalev et al. 2022).

The observed heterozygosity (H_o_) was similar between adults and juveniles in the natural regeneration across all sites. This may indicate that the frequency of heterozygotes on each site has not been reduced suggesting an efficient transfer of alleles to the next generation of trees. These results follow the observations of Liesebach et al. (2024) in *Picea abies* and *Fagus sylvatica* stands in Europe, where H_o_ was consistent between adults and natural regeneration; however, the authors reported differences in the effective population size N_e_.

The N_e_ is a genetic diversity estimator that is gaining importance due to its recent inclusion in the Convention on Biological Diversity (CBD) Global Biodiversity Framework (Frankham 2022; Heuertz et al. 2023; Hoban et al. 2020, 2023). None of the sites had an N_e_ > 500, which is the current target established to ensure sufficient maintenance of genetic variation in the long-term (Hoban et al. 2020). Only Stourhead1 juveniles (N_e_ = 97.8) is close to N_e_ = 100 which is considered the minimum N_e_ to avoid short-term inbreeding depression (Frankham et al. 2014). We estimated the N_e_ using only monomorphic SNPs but these were fewer than the 180 minimum SNPs recommended (Waples and Do 2010), thus, our results must be interpreted with caution; however, we ensured no linkage among the study SNPs, which is one of the main assumptions of the method (Luikart et al. 2010; Waples et al. 2016; Waples and Do 2010). Although the N_e_ estimated in the species’ natural range by Shalev et al. (2022) is higher than our results (N_e_ = 270.3), it also revealed low levels of genetic diversity across the natural distribution which follows the low heterozygosity levels reported.

On average across all the study woodlands, the proportion of assigned parents-offspring relationships detected across sites was 26.25% at 95% confidence indicating a majority of parents being either from individuals located within the sites but not sampled or from outside the study sites. Slightly lower levels of assignment were found in Duminil et al. (2016), who investigated the parental assignment on a high selfing tree species (*Baillonella toxisperma*) and found only 10% of the offspring could be assigned to a sampled parent. Here, the lowest assignment level was at Longleat1 (11%), which is the only site that is isolated by > 2 Km from other *T. plicata* stands (Table 1), suggesting potential pollen flow from more remote stands. It also had the highest number of offspring related to one single parent, indicating the potential for a relatively high level of half-sibship. Bagley Wood gave the highest proportion of assignments (37%) and a low number of offspring assigned to one single parent, suggesting a lower level of half-sibships. Bagley Wood is the smallest site in terms of area but had the highest density of *T. plicata* trees (Table 1), which could explain the high number of assigned trees. Stourhead1 had 35% of assigned parents, indicating that the patchy distribution of these sites (Table 1) does not prevent gene flow among the patches. This is also demonstrated by looking at the distance between individuals from separated patches being assigned (Stourhead1 = 310.4 m and Bagley Wood = 1,456.0 m) (Table 6). The high rate of selfing reported in *T. plicata* (O’Connell et al. 2001) may also have affected the proportion of assigned parent-offspring pairs found in our study.

### 5. Practical application and conclusions

We developed and validated a set of markers (SNP set2) to evaluate genetic variation and genetic diversity in *T. plicata* populations. These SNP markers were carefully selected and could be useful in genetic investigations of *T. plicata* populations, offering a reliable, cost-effective, and time-efficient method of testing many samples. We found many monomorphic SNPs unique to the adults from Longleat1, indicative of a lower genetic variation within this site and a distinct genetic background. When the private monomorphic SNPs were excluded, the genetic differentiation was low between Bagley Wood and Stourhead1, suggesting that the stands were made up of materials of similar origin. We found consistent heterozygosity levels in adults and juveniles across all sites, indicating that the natural regeneration may hold similar genetic diversity and gene pool size than the parents. This observation suggests that the potential for genetic adaptation to changing conditions including climate warming is maintained across generations. However, the effective population size was low to very low, with only Stourhead1 juveniles observed to have N_e_ ≈ 100. The low N_e_ suggests that the genetic makeup of *T. plicata* plantations in the UK may be rather narrow and potentially need an injection of diversity to ensure their genetic viability in the long-term. We found that the stand (Stourhead2) with a more advanced transformation toward CCF as indicated by species and structural diversity had very low N_e_ levels, in contrast to other stands (e.g. Bagley Wood) which are less diverse but have higher heterozygosity and N_e_ levels. This observation suggests that silviculture treatments such as stand thinnings to encourage species diversity may have negative consequences for N_e_ and genetic diversity. This trade-off means that forests we perceive as more diverse or naturalised may not hold enough standing genetic diversity to face current and future disturbances caused by climate change.

Our study combined analyses of CCF transformation and genetic diversity in four *T. plicata* populations in the UK thus producing integrated insights for woodland management. Future studies may aim to improve our understanding of the species as a non-native, e.g. by mapping the origin of UK planted materials to areas in its natural range to uncover the population genetics of monomorphic alleles only observed in Longleat1 adults. A better understanding of variations in N_e_ would also be of interest both in the natural range and in areas where it has been introduced with implications for genetic resource management. Our results suggest that a potential injection of genetic diversity (e.g. through planting) may be needed in woodlands under selective harvesting in order to maintain or increase genetic diversity and N_e_.

## Supporting information

Online resource

## Acknowledgements

The authors acknowledge the forestry estates where the samples came from: Stourhead (Western), Longleat (Wiltshire) and Bagley Wood. We also thank Barley Rose Collier Harris for helping with sampling. We want to thank Will Hoare, George MacKay and Emily Farley for their assistance in both laboratory and field work. The authors would like to acknowledge the use of the University of Oxford Advanced Research Computing (ARC) facility in carrying out this work. http://dx.doi.org/10.5281/zenodo.22558

## Declarations

### Authors’ contribution statements

LG drafted the manuscript and both LG and JM revised the main manuscript. LG conceived and planned the experiments, performed laboratory and statistical analyses and prepared the figures. LG and EG performed the sampling and developed the field survey. JM supervised the project.

## Data Archiving Statement

The raw NGS read data was deposited to NCBI’s Short Read Archive (SRA). The BioProject ID is PRJNA1081148 and the SRA accession numbers range from SAMN40149401 to SAMN40149480. SNP set1 and SNP set2 information are uploaded to Dryad (doi:xxxxxxx).

