## Supplementary material for "Low effective population size but high heterozygosity revealed by SNP analyses in adult and juvenile *Thuja plicata* in UK Woodlands": Online resource

Tree Genetics and Genomes

### Supplementary Information

Online resource 1. Number of fragments recovered by simulating the genome digestion using the potential enzymes.

| Enzyme | Fragment size 126-300 bp |
|---|---|
| ApeKI | 565,079 |
| BfaI | 7,962,110 |
| PstI_MspI | 112,191 |
| NsiI_MspI | 319,908 |
| SbfI_MspI | 8,236 |

Online resource 2. Parameters name and value used in Ipyrad pipeline

| Parameter name | value |
|---|---|
| Assembly method | reference |
| Reference sequence | pairddrad |
| Restriction overhang | TGCAG,CG |
| Max low quality bases | 5 |
| Phred Qscore offset | 43 |
| Min depth statistical | 6 |
| Min depth majrule | 6 |
| Max depth | 10000 |
| Cluster threshold | 0.90 |
| Max barcode mismatch | 0 |
| Filter adapters | 2 |

| | |
|---|---|
| Filter min trim len | 35 |
| Max alleles consens | 2 |
| Max Ns consens | 0.05 |
| Max Hs consens | 0.05 |
| Min samples locus | 4 |
| Max SNPs locus | 0.2 |
| Max Indels locus | 8 |
| Max shared Hs locus | 0.5 |
| Trim reads | 0, 0, 0, 0 |
| Trim loci | 0, 0, 0, 0 |

Online Resource 3. Final summary of statistics of the Ipyrad assembly run per sample. Accession SRA number (SRA). Reads raw: number of raw reads demultiplexed for each sample, Reads passed filter: number of reads passed filter based on quality scores, Refseq mapped reads: number of reads mapped to the reference genome, Refseq unmapped reads: number of reads that did not mapped to the reference genome, Clusters total: Total clusters found, Clusters hidepth: number of clusters that pass the mindepth thresholds, Hetero est: heterozygosity estimation, Error est: Sequencing error rate, Reads consens: number of final reads after error filtering and Loci in assembly: number of loci recovered per sample.

| SampleID | Reads raw | Reads passed filter | Refseq mapped reads | Refseq unmapped reads | Clusters total | Clusters hidepth | Hetero est | Error est | Reads consens | Loci in assembly |
|---|---|---|---|---|---|---|---|---|---|---|
| B1PA16 | 18456767 | 18159596 | 17515113 | 644483 | 104294 | 55355 | 0.007844 | 0.000187 | 24855 | 21154 |
| B1PA17 | 19537438 | 19223586 | 18637723 | 585863 | 74214 | 41425 | 0.00232 | 0.00019 | 21986 | 19379 |
| B1PA26 | 28719642 | 28244069 | 26843172 | 1400897 | 156332 | 80851 | 0.002458 | 0.000154 | 28959 | 23162 |
| B1PA3 | 25120931 | 24719238 | 23853196 | 866042 | 115337 | 58625 | 0.002632 | 0.000155 | 25615 | 21778 |
| B1PA33 | 4271424 | 4192103 | 3906045 | 286058 | 61659 | 29361 | 0.00222 | 0.000489 | 15902 | 14543 |
| B1PA35 | 6716150 | 6598027 | 6332132 | 265895 | 58435 | 28958 | 0.002604 | 0.000423 | 17651 | 15856 |
| B1PA37 | 17692561 | 17370021 | 16777371 | 592650 | 86969 | 46385 | 0.002109 | 0.000252 | 21590 | 19018 |
| B1PA45 | 24266675 | 23840745 | 23196252 | 644493 | 80547 | 45430 | 0.002645 | 0.000168 | 24998 | 21195 |
| B1PA52 | 23574560 | 23159694 | 21962088 | 1197606 | 174182 | 85330 | 0.002387 | 0.000204 | 27087 | 22236 |
| B1PR106 | 17990112 | 17695293 | 17054392 | 640901 | 92342 | 48204 | 0.002945 | 0.000166 | 22155 | 19065 |
| B1PR109 | 19286705 | 19009397 | 18457941 | 551456 | 69569 | 41041 | 0.002565 | 0.000189 | 23261 | 20228 |
| B1PR135 | 7647035 | 7491046 | 7270352 | 220694 | 37373 | 22359 | 0.002909 | 0.000489 | 17403 | 15562 |
| B1PR142 | 5877271 | 5768749 | 5609080 | 159669 | 34349 | 21602 | 0.002651 | 0.000452 | 18196 | 16137 |
| B1PR35 | 31891988 | 31345061 | 30209781 | 1135280 | 146590 | 73539 | 0.002682 | 0.00017 | 28682 | 22940 |
| B1PR52 | 26661277 | 26222864 | 25315304 | 907560 | 98487 | 52388 | 0.002458 | 0.000154 | 26621 | 21773 |
| B1PR59 | 17072105 | 16828577 | 16411793 | 416784 | 65088 | 34944 | 0.007935 | 0.000235 | 24319 | 20150 |
| B1PR78 | 18568664 | 18303819 | 17077403 | 1226416 | 162838 | 77873 | 0.001835 | 0.000205 | 21128 | 17909 |
| B1PR92 | 6575160 | 6460195 | 6232016 | 228179 | 37887 | 22656 | 0.002692 | 0.000438 | 17297 | 15542 |
| B1PR96 | 14025729 | 13803804 | 13449098 | 354706 | 44235 | 29046 | 0.002229 | 0.000249 | 21222 | 18665 |
| L1PA1 | 2412254 | 2373280 | 2056446 | 316834 | 67128 | 28803 | 0.001986 | 0.000467 | 14211 | 13052 |
| L1PA1b | 23906361 | 23535822 | 20495243 | 3040579 | 265382 | 128505 | 0.001765 | 0.000187 | 25119 | 19700 |
| L1PA21 | 30201716 | 29637923 | 27767279 | 1870644 | 158627 | 78742 | 0.007844 | 0.000187 | 26399 | 21439 |
| L1PA22 | 39012765 | 38410828 | 30384598 | 8026230 | 501110 | 250448 | 0.001662 | 0.000233 | 31668 | 20970 |

| SampleID | Reads raw | Reads passed filter | Refseq mapped reads | Refseq unmapped reads | Clusters total | Clusters hidepth | Hetero est | Error est | Reads consens | Loci in assembly |
|---|---|---|---|---|---|---|---|---|---|---|
| L1PA39 | 17677918 | 17369674 | 15448261 | 1921413 | 237883 | 113294 | 0.006437 | 0.000475 | 23385 | 18931 |
| L1PA41 | 19134922 | 18802077 | 16217219 | 2584858 | 204633 | 94774 | 0.001909 | 0.000407 | 23064 | 19137 |
| L1PA42 | 28246955 | 27788324 | 20862661 | 6925663 | 393548 | 192749 | 0.001983 | 0.000303 | 29769 | 21323 |
| L1PA7 | 12670635 | 12438997 | 11490295 | 948702 | 121892 | 59835 | 0.006312 | 0.000625 | 21586 | 18531 |
| L1PA8 | 6209772 | 6078319 | 4503369 | 1574950 | 153593 | 62983 | 0.002035 | 0.000475 | 16536 | 14597 |
| L1PR109 | 17335906 | 17050214 | 15568784 | 1481430 | 122685 | 61607 | 0.006796 | 0.000239 | 22071 | 18885 |
| L1PR121 | 2656813 | 2604449 | 2507573 | 96876 | 24775 | 16587 | 0.002279 | 0.000534 | 14088 | 12868 |
| L1PR19 | 20536022 | 20182869 | 17490884 | 2691985 | 254434 | 117766 | 0.002166 | 0.000298 | 26231 | 20887 |
| L1PR35 | 30849686 | 30251526 | 29205931 | 1045595 | 90113 | 50158 | 0.002429 | 0.000181 | 26741 | 21787 |
| L1PR42 | 10345995 | 10168553 | 9674771 | 493782 | 61262 | 33094 | 0.002268 | 0.000563 | 18788 | 16758 |
| L1PR49 | 4662460 | 4561428 | 4047368 | 514060 | 66770 | 31223 | 0.002195 | 0.000626 | 16240 | 14301 |
| L1PR56 | 41737756 | 40933375 | 25899321 | 15034054 | 703355 | 343379 | 0.002073 | 0.000286 | 37026 | 20280 |
| L1PR87 | 14384768 | 14155605 | 13504166 | 651439 | 69035 | 37976 | 0.002046 | 0.000357 | 20043 | 17659 |
| L1PR88 | 5803441 | 5713306 | 5134337 | 578969 | 89960 | 41148 | 0.002706 | 0.000597 | 17857 | 16106 |
| S1PA2 | 14730633 | 14480400 | 12768272 | 1712128 | 123015 | 63660 | 0.002724 | 0.000477 | 23585 | 19845 |
| S1PA24 | 15875952 | 15620984 | 14920390 | 700594 | 81087 | 44884 | 0.001958 | 0.000237 | 21540 | 18532 |
| S1PA26 | 15749993 | 15490544 | 15117856 | 372688 | 49424 | 31152 | 0.002387 | 0.000307 | 23171 | 19724 |
| S1PA31 | 15230305 | 14969584 | 14599620 | 369964 | 49725 | 30977 | 0.003032 | 0.000377 | 22745 | 19627 |
| S1PA36 | 21879133 | 21538217 | 21034287 | 503930 | 66716 | 39344 | 0.002725 | 0.00031 | 26445 | 21798 |
| S1PA37 | 18909594 | 18615027 | 18207444 | 407583 | 51199 | 31747 | 0.007844 | 0.000187 | 24174 | 20175 |
| S1PA39 | 9267306 | 9101483 | 8807791 | 293692 | 47059 | 26326 | 0.002502 | 0.000475 | 18886 | 16705 |
| S1PA44 | 2326566 | 2269205 | 2171755 | 97450 | 25530 | 17368 | 0.002129 | 0.000501 | 15333 | 13714 |
| S1PA46 | 2268813 | 2229595 | 2133191 | 96404 | 25106 | 16849 | 0.002478 | 0.000517 | 14121 | 12997 |
| S1PA50 | 14990288 | 14757917 | 14026688 | 731229 | 79870 | 44086 | 0.00202 | 0.000392 | 19750 | 17218 |
| S1PR101 | 19852831 | 19556414 | 18580276 | 976138 | 120189 | 60282 | 0.00331 | 0.00027 | 25108 | 21128 |
| S1PR11 | 6636533 | 6525054 | 6352926 | 172128 | 38017 | 25533 | 0.0032 | 0.000771 | 19800 | 17585 |
| S1PR135 | 5032566 | 4949922 | 3978431 | 971491 | 106795 | 46611 | 0.002405 | 0.000397 | 16174 | 14529 |
| S1PR167 | 16106380 | 15819117 | 15473566 | 345551 | 46363 | 30349 | 0.003114 | 0.00028 | 23515 | 20108 |
| S1PR172 | 23853209 | 23452636 | 22913318 | 539318 | 55948 | 36680 | 0.002544 | 0.000197 | 28269 | 23245 |
| S1PR32 | 29744043 | 29290876 | 26259610 | 3031266 | 211572 | 103585 | 0.002422 | 0.000216 | 27346 | 21997 |
| S1PR56 | 33989129 | 33437259 | 31945966 | 1491293 | 119509 | 65475 | 0.002682 | 0.00017 | 29109 | 23559 |
| S1PR58 | 17511511 | 17192210 | 16641665 | 550545 | 63253 | 37268 | 0.009565 | 0.000256 | 23574 | 19951 |
| S1PR82 | 6770413 | 6662271 | 6461296 | 200975 | 41827 | 26698 | 0.002584 | 0.000523 | 19201 | 17172 |
| S4PA10 | 11461965 | 11287605 | 11001190 | 286415 | 49520 | 29274 | 0.00247 | 0.000529 | 18808 | 16509 |
| S4PA13 | 12825847 | 12608168 | 12014037 | 594131 | 82998 | 43300 | 0.002376 | 0.00039 | 20037 | 17617 |
| S4PA15 | 13147178 | 12967756 | 12486692 | 481064 | 82658 | 40308 | 0.002236 | 0.000343 | 19650 | 17192 |
| S4PA19 | 24376469 | 23966060 | 23223028 | 743032 | 74549 | 41222 | 0.002199 | 0.000199 | 22305 | 19308 |
| S4PA20 | 18205817 | 17919781 | 17124914 | 794867 | 106849 | 56087 | 0.002294 | 0.0003 | 21138 | 18210 |
| S4PA21 | 9418500 | 9290060 | 8994494 | 295566 | 55394 | 28405 | 0.00187 | 0.000406 | 17511 | 15689 |
| S4PA25 | 14498193 | 14287536 | 13795052 | 492484 | 63040 | 35967 | 0.006906 | 0.000362 | 21844 | 18958 |
| S4PA26 | 16420977 | 16177287 | 15614581 | 562706 | 67147 | 37456 | 0.001406 | 0.000283 | 21738 | 18709 |
| S4PA3 | 8182460 | 8036503 | 7609384 | 427119 | 61057 | 34424 | 0.002889 | 0.000943 | 18606 | 16362 |
| S4PA35 | 7502708 | 7359091 | 6796538 | 562553 | 66266 | 32027 | 0.006312 | 0.000625 | 17433 | 15293 |
| S4PR35 | 2620308 | 2556327 | 2456159 | 100168 | 27314 | 18067 | 0.002655 | 0.000572 | 15621 | 13694 |
| S4PR42 | 6163660 | 6037215 | 5875635 | 161580 | 32959 | 21088 | 0.002805 | 0.000596 | 17983 | 15864 |
| S4PR47 | 9331681 | 9167501 | 8272593 | 894908 | 89289 | 43577 | 0.002919 | 0.000757 | 18760 | 16294 |
| S4PR77 | 20247411 | 19894843 | 19348648 | 546195 | 69211 | 38244 | 0.002078 | 0.00033 | 21702 | 18612 |
| S4PR81 | 17159707 | 16837509 | 16321420 | 516089 | 75501 | 39781 | 0.001983 | 0.000303 | 22697 | 19299 |
| S4PR89 | 23326493 | 22974434 | 22061128 | 913306 | 111321 | 55644 | 0.00105 | 0.000171 | 21896 | 18030 |
| S4PR90 | 24711091 | 24296552 | 23811475 | 485077 | 52411 | 32906 | 0.007844 | 0.000187 | 24094 | 19943 |
| S4PR96 | 15996841 | 15721257 | 15322535 | 398722 | 57627 | 32872 | 0.002522 | 0.000339 | 21798 | 18367 |
| S4PR96b | 19355832 | 19048090 | 18602150 | 445940 | 63978 | 36894 | 0.002591 | 0.000246 | 23532 | 19281 |

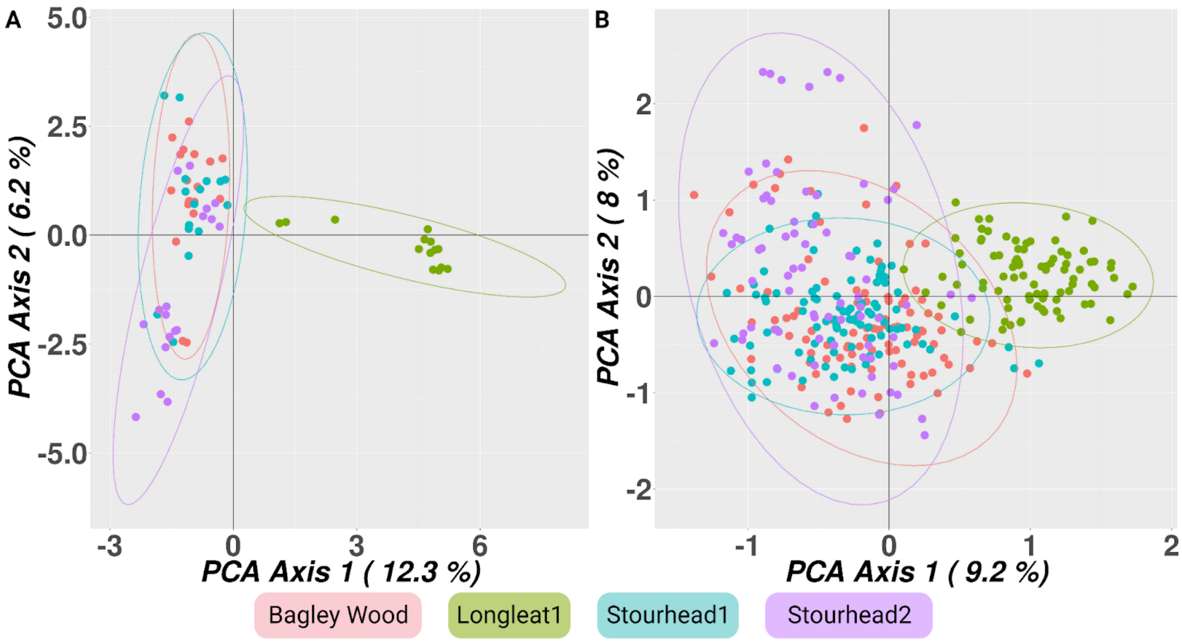


Online Resource 4. Analysis of principal Coordinates (PCoA) results using A) SNP set1 and B) SNP set2 across the four study sites.
